# «Studies *in vitro* e *in vivo* of Phage Therapy Medical Products (PTMPs) targeting clinical strains of *Klebsiella pneumoniae* belonging to the clone ST512»

**DOI:** 10.1101/2024.11.20.624559

**Authors:** Inés Bleriot, Lucía Blasco, Patricia Fernández-Grela, Laura Fernández-García, Lucia Armán, Clara Ibarguren, Concha Ortiz-Cartagena, Antonio Barrio-Pujante, José Ramón Paño, Jesús Oteo-Iglesias, María Tomás

## Abstract

The widespread incidence of antimicrobial resistance has created renewed interest in the use of alternative antimicrobial treatments such as phage therapy. Phages are viruses that infect bacteria and generally have a narrow bacteria host-range. Combining phages with antibiotics can prevent the emergence of bacterial resistance. The aim of the present study was to develop phage therapy medical products (PTMPs) targeting clinical isolates of carbapenems-producing *Klebsiella pneumoniae* belonging to the high-risk clone ST512. From a collection of twenty-two seed of lytic phages sequenced belonging to MePRAM collection, four were used to generate PTMPs (CAC_Kpn1 and CAC_Kpn2). These PTMPs were partly active against three of the clinical strains of clone ST512 (A, B and C). The use of Appelmans method in the CAC_Kpn1_ad (adapted CAC_Kpn1) yielded a significant increase in the efficacy against strain A, while adapted CAC_Kpn2 (CAC_Kpn2_ad) only effectively reduced bacterial survival when combined with ½ x MIC ß-lactam antibiotic meropenem for 24 h in clinical strains B and C, showed after this time, resistance to PTMPs. In addition, the amounts of endotoxin released by the PTMPs were quantified and subsequently reduced in preparation for *in vivo* use of the PTMPs in *Galleria mellonella* infection model confirming the *in vitro* results from the CAC_Kpn1_ad and CAC_Kpn2_ad.

## INTRODUCTION

The current widespread incidence of antimicrobial resistance (AMR) is a global health problem. Today, an estimated 700,000 people lose their lives as a result of antimicrobial resistance; if left unaddressed, this problem is expected to lead to an estimated 10 million deaths per year by 2050 (1). The lack of treatments for multidrug-resistant (MDR) bacteria has placed the spotlight on alternative therapies such as bacteriophages (phages), which fell out of favour in the West after the appearance of antibiotics. Phages are viruses that infect bacteria, generally within a narrow bacteria host-range, and are found in all habitats where bacteria are present (2). Phages thus having certain advantages over the use of conventional antibiotics, notably high host specificity and low cost (3). However, as phages and bacteria are constant co-evolving, the bacterial hosts can become resistant to the phages that infect them. This is a major problem that can lead to the failure of phage therapy (4, 5). One option is to develop phage therapy medical products (PTMPs) to prevent phage resistance (6, 7), in an approach that has been successfully used in the Elviana Institute of Bacteriophage, Microbiology and Virology in Georgia. PTMPs are generated by mixing several phages together to enhance their effect against MDR bacterial infections. However, it is not always possible to isolate suitable seed of phages or design PTMPs that target the most important clinical isolates. The Appelmans protocol, also known as phage training, is one of the approaches used to solve this problem (8–10). This technique has been successfully used for decades in Georgia and has recently been revived in the West. The technique consists of the successive passage of a PTMPs through a series of bacterial strains of interest, most of which are initially phage refractory, until the bacteria finally undergo lysis (10, 11). Another separate or complementary technique is to combine the PTMPs with antibiotics to improve the efficacy of the antimicrobial against MDR bacteria (12–14). The combined products can have different types of effects on each other: antagonistic, synergistic or no effect. In this case, the effect of interest is synergism, whereby the combination of two agents used have a greater effect than the sum of the effects caused by each agent separately (15). The term phage-antibiotic synergy (PAS) usually refers to the use of sub-lethal concentrations of antibiotics, generally ß-lactam and quinolones, with phages (12, 13, 16–18).

In this study, the pathogen of interest was the MDR bacteria *Klebsiella pneumoniae*, against which novel treatments are urgently required. *K. pneumoniae* is an encapsulated, rod-shaped, non-motile, lactose-fermenting, encapsulated bacillus that is highly ubiquitous, being found in the environment as well as in the microbiota of healthy individuals. However, it is also an opportunistic pathogen that can cause a wide range of infections, including pneumonia, soft tissue infections, surgical wound infections, urinary tract infections and sepsis. This pathogen has provoked great interest due to the increase in the incidence of severe infections that it causes and also the scarcity of available treatments (19). The number of hospitals affected by outbreaks of infections caused by *K. pneumoniae* has increased in recent years (20, 21). The carbapenemase-producing *K. pneumoniae* clones of sequence types ST258 and ST512 are currently the dominant clones by which *K. pneumoniae* has spread worldwide (22–25).

The aim of the present study was to create adapted PTMPs that can be used to target clinical three isolates of carbapenemase-producing *K. pneumoniae* belonging to the high-risk clone ST512.

## RESULTS

### Host-range assay

Four seed of phages with a wide host-range, vB_KpnM-VAC13 (12.77 %), vB_KpnP-VAC25 (25.53 %), vB_KpnM-VAC36 (19.14 %) and vB_KpnM-VAC66 (6.38 %), were selected for generating two PTMPs against *K. pneumoniae* strains A, B and C (Figure 1). The CAC_Kpn1 PTMP, effective against strain A, was composed of seed of phages vB_KpnM-VAC13, vB_KpnM-VAC36 and vB_KpnM-VAC66, while the CAC_Kpn2 PTMP, effective against strains B and C, was composed of seed of phages vB_KpnP-VAC25, vB_KpnM-VAC36, and vB_KpnM-VAC66. Comparative genomic analysis was conducted to establish the relatedness of the selected seed of phages (Figure 2). The comparison showed that seed of phages vB_KpnM-VAC13 and vB_KpnM-VAC66 are very similar (Query cover: 95 % and Identity: 97.56 %) and that seed of phages vB_KpnM-VAC13, vB_KpnM-VAC66 and vB_KpnM-VAC36 are different (vB_KpnM-VAC13 vs vB_KpnM-VAC36: Query cover 0% and Identity: 80.80 %; vB_KpnM-VAC66 vs vB_KpnM-36: Query cover: 3 % and Identity: 85.05 %). Phages vB_KpnM-VAC13, vB_KpnM-VAC36 and vB_KpnM-VAC66 belong respectively to the genera *Slopekvirus* and *Marfavirus* (Figure 2). However, phage vB_KpnP-VAC25, which belongs to the genus *Drulisvirus* (Figure 2), is very different from the other three phages (the comparison of all phages vs vB_KpnP-VAC25 did not reveal significant similarity).

**Figure 1.**
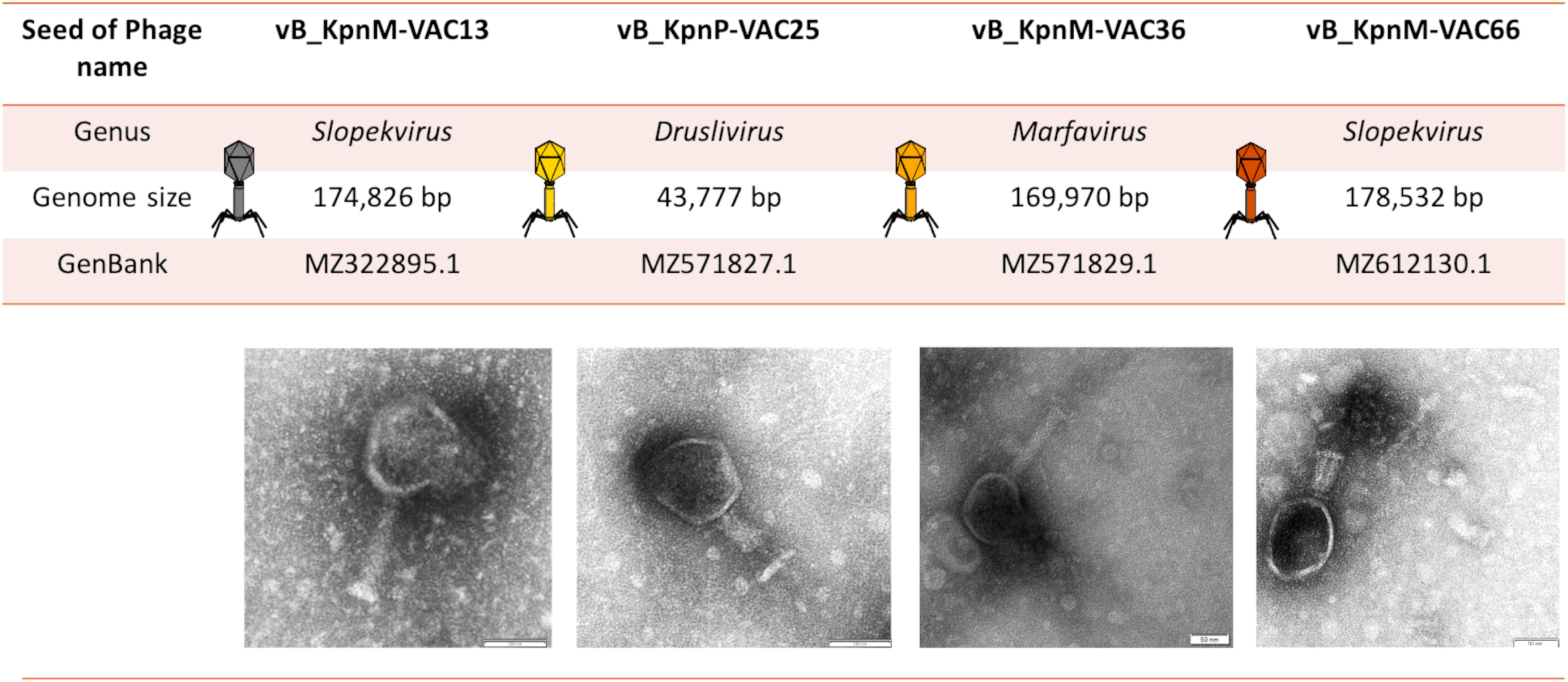
Morphological characteristics of the selected seed of phages. The morphology of vB_KpnM-VAC13, vB_KpnM-VAC36 and vB_KpnM-66 is typical of myoviruses, while the morphology of vB_KpnP-VAC25 is typical of a podovirus.

**Figure 2.**
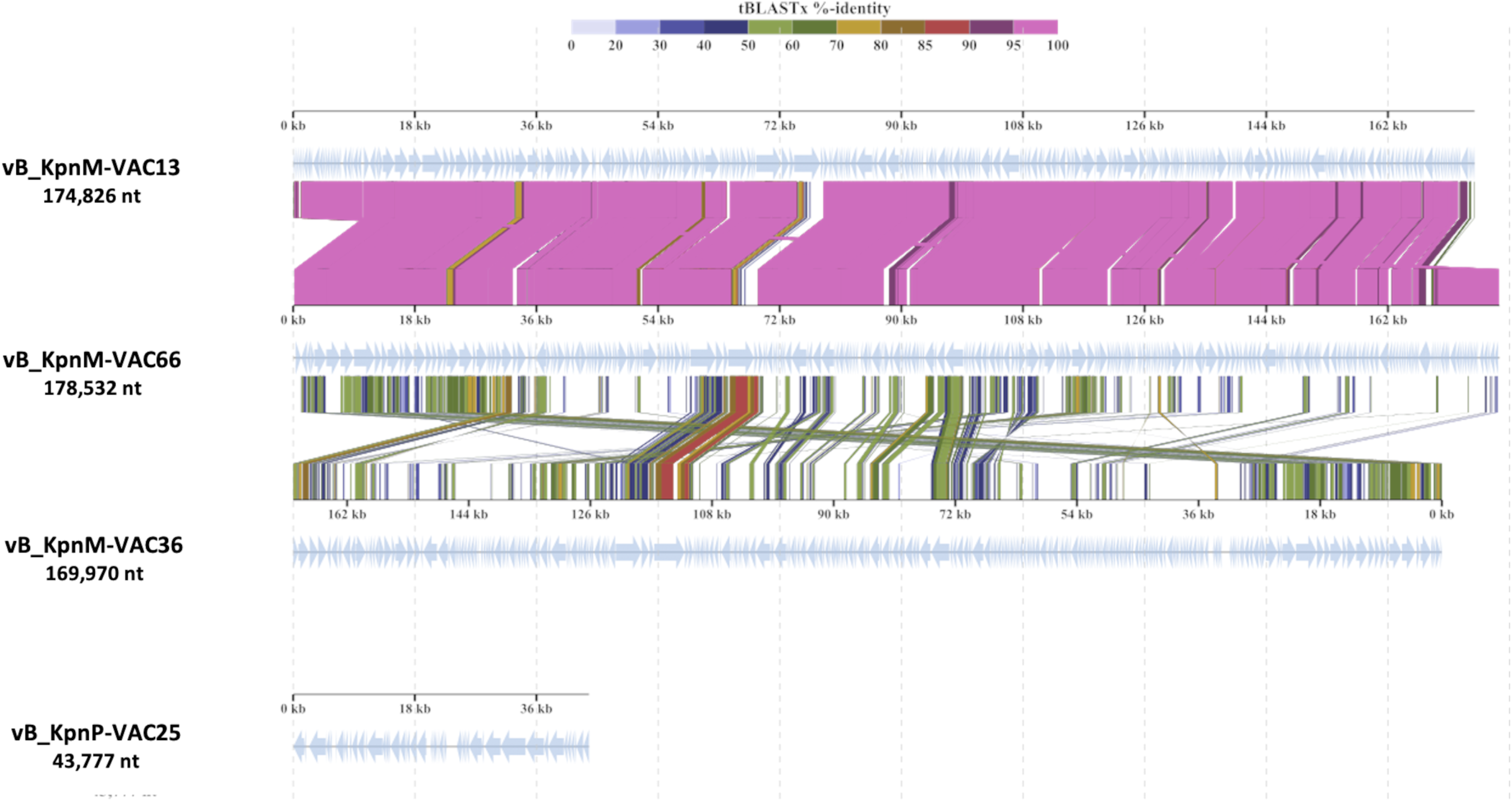
Graphic comparison of the homology of the four seed of phages selected to elaborate PTMPs CAC_Kpn1 and CAC_Kpn2. The schematic representation was constructed with the VipTree server (https://www.genome.jp/viptree/).

### Electron microscopic analysis of seed of phages

TEM analysis revealed the morphology of the four seed of phages selected. The morphology of vB_KpnM-13, vB_KpnM-36 and vB_KpnM-VAC66 was typical of myoviruses, while that of vB_KpnP-VAC25 was characteristic of a podovirus (Figure 1).

### Infectivity assay in liquid media

The infectivity assay in liquid medium performed with each of the seed of phages (Figure 3) revealed that all (vB_KpnM-VAC13, vB_KpnM-VAC36 and vB_KpnM-VAC66) were highly infective in strain A in liquid medium, with seed of phages vB_KpnM-VAC36 and vB_KpnM-VAC66 being the most infective (Figure 3 A). For strains B and C, seed of phages vB_KpnM-VAC36 and vB_KpnM-VAC66 produced a mild infection followed by the appearance of resistance (Figure 3 B and C). The PTMP CAC_Kpn1, which infects strain A, showed an efficacy of 100 % during the first 7 h, which resulted in complete inhibition of bacterial growth (Figure 4 A). However, resistance emerged after 7 h. These results were confirmed by the CFU/mL and PFU/mL counts (Table S1). Briefly, a significant decrease in the CFU/mL count was observed after 12 h (p-value < 0.05), resulting in a slight increase in the PFU/mL; this shows that the PTMPs are effective against strain A; however, after 22 h significant increases (p-value < 0.05) were observed in the CFU/mL and PFU/mL counts, corresponding to the emergence of resistance observed in the absorbance curve. In the case of the PTMP CAC_Kpn2 for strain B, the CFU/mL counts significant increased after 12 h (p-value < 0.05) and thereafter continued to increase until the end of the experiment, at 22 h (Figure 4 B). The same phenomen was observed with the PTMP CAC_Kpn2 for the strain C, with a significant increase at 12 h (p-value < 0.05) and a slight increase at 22 h (Figure 4 C).

**Figure 3.**
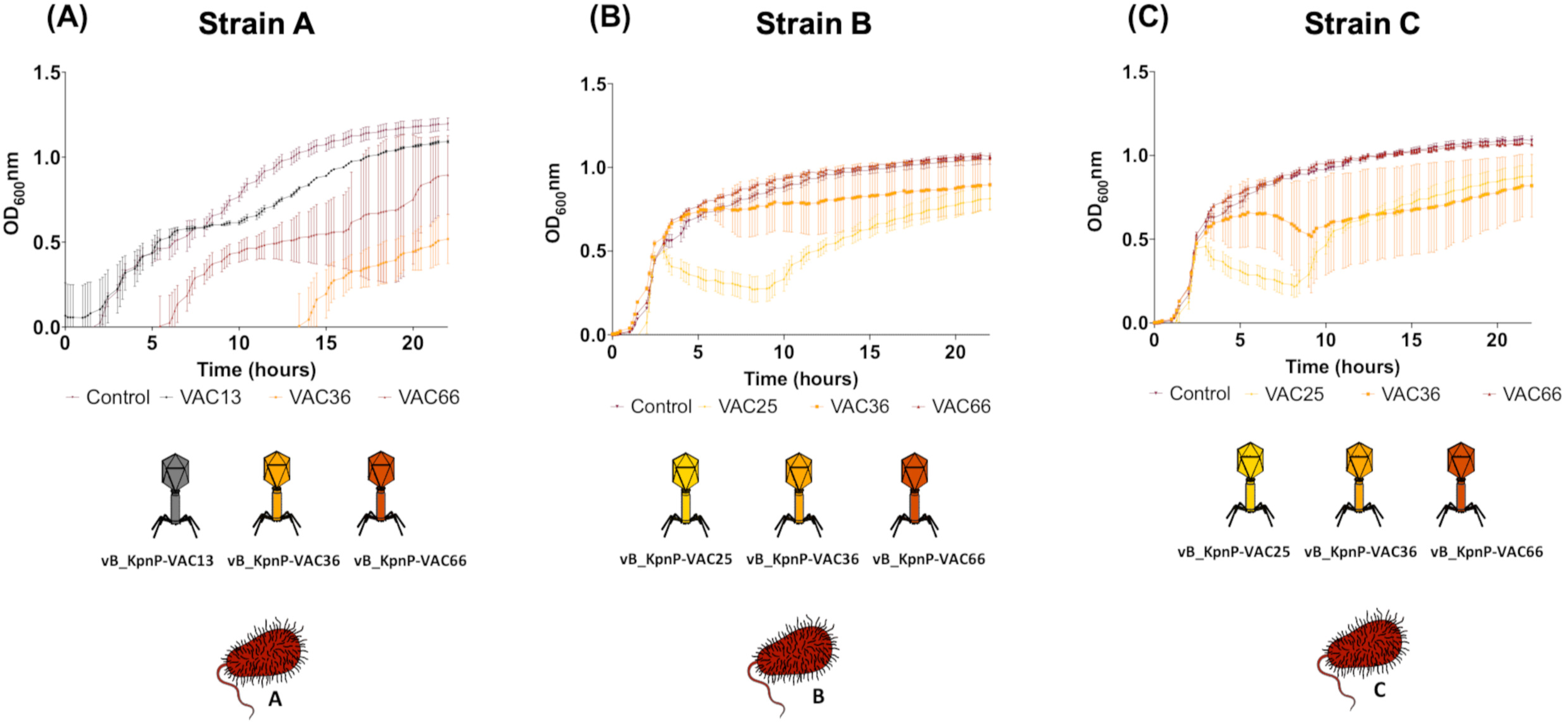
Infection curves for strains **(A)** A, **(B)** B and **(C)** C with individual seed of phages at 24 h.

**Figure 4.**
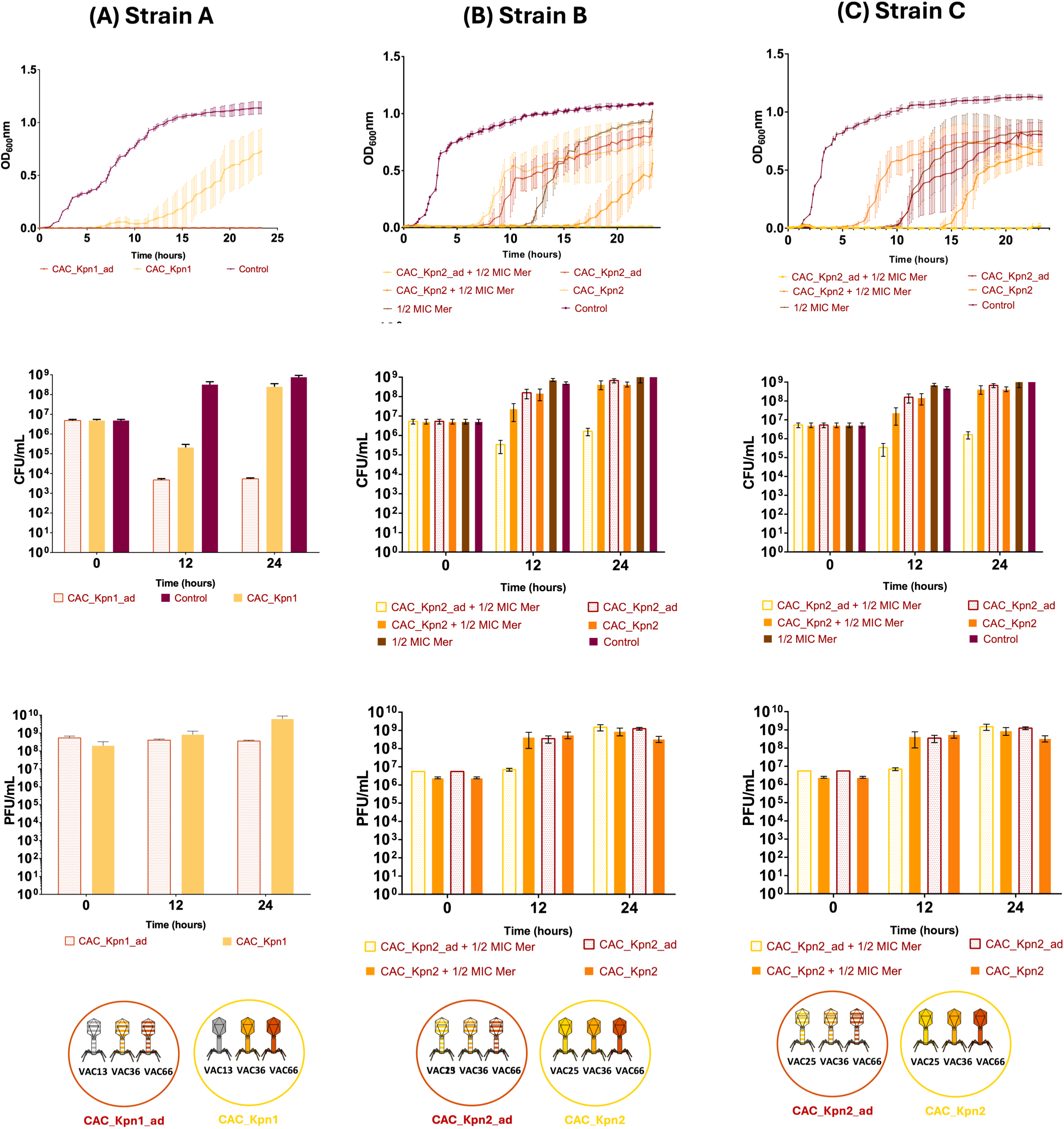
Infection curves for strains **(A)** A, **(B)** B and **(C)** C with adapted and non-adapted PTMPs. The curves for the adapted and no-adapted PTMPs in the presence of ½ MIC of meropenem for strains B and C are also shown. The CFU/mL and PFU/mL counts are shown for all strains.

### Phage training: Adaptation of the PTMPs

In line with the results obtained in the liquid test of the CAC_Kpn1 and CAC_Kpn2 PTMPs, both were adapted using Appelmans protocol, with the aim of improving the therapeutic efficacy and reducing the emergence of resistance. Adaptation of the CAC_Kpn1 PTMP (CAC_Kpn1_ad), which infects strain A, greatly improved the infectivity, yielding 100 % efficacy over 22 h. These results were confirmed by CFU/mL and PFU/mL counts (Table S1), which revealed a significant decrease in CFU/mL count (p-value < 0.05) and maintenance of the PFU/mL count (Figure 4 A). On the other hand, in the case of the CAC_Kpn2_ad PTMP, the CFU/mL count increased over time, as well as an increase in PFU/mL (Figure 4 B and C; Table S1). As the efficacy of the adapted PTMPs was not very clear, further trials were carried out by adding a ß-lactam antibiotic (meropenem) to the adapted PTMPs to reduce the potential emergence of resistance over time.

### Meropenem minimal inhibitory concentration assay and infectivity assay in liquid media with ½ MIC of meropenem

A microdilution broth assay was performed to determine the meropenem MIC for clinical isolates of *K. pneumoniae* B and C, yielding values of 16 mg/L and 8 mg/L, respectively. The liquid infectivity test performed using the combination of ½ of MIC of meropenem and the PTMPs CAC_Kpn2_ad revealed a great improvement in efficacy, reaching 100 % within 20 h for strain B and within 22 h (Figure 4 B) in for strain C (Figure 4 C). Regarding the CFU/mL counts for strains B and C, a significant decrease in CFU/mL counts was observed at 12 h (P-value < 0.05). The PFU/mL counts increased over time in both strains (Table S1).. Therefore, PAS was obtained.

### Determination of the rate of emergence of bacterial mutants

Determination of the rate of emergence of phage-resistant mutant bacterial cells (Table 1), revealed that the strain A with the CAC_Kpn1_ad show a significant lower value in comparison with the other two PTMPs (p-value < 0.05). For the strain B, the results revealed that the combination of PTMP and antibiotic had a significantly lower effect than the monotherapy (p-value < 0.05) and together these yielded a synergic effect. On the other hand, for strain C, there was a significant difference between the CAC_Kpn2_ad and the CAC_Kpn2_ad in combination with ½ MIC of meropenem (p-value < 0.05); however, there was no significant difference between ½ MIC of meropenem and the CAC_Kpn2_ad in combination with ½ MIC of meropenem (p-value = 0.1134). Significant differences were observed in the efficacy of the adapted PTMPs between the strains B and C (p-value < 0.05).

**Table 1.** Frequency of occurrence of bacterial resistance of strain A, B and C. The results are expressed as frequency of mutations per bacterial cell.

|  | Strain A | Strain B | Strain C |
| --- | --- | --- | --- |
| *Adapted PTMPs* | $2.19 \times 10^{-7} \pm 9.15 \times 10^{-8}$ | $1.26 \times 10^{-4} \pm 6.71 \times 10^{-5}$ | $4.24 \times 10^{-2} \pm 6.03 \times 10^{-3}$ |
| $\frac{1}{2}$ MIC Mer | | $2.70 \times 10^{-4} \pm 3.19 \times 10^{-5}$ | $4.95 \times 10^{-4} \pm 9.99 \times 10^{-5}$ |
| *Adapted PTPMs + $\frac{1}{2}$ MIC Mer* | | $1.75 \times 10^{-5} \pm 7.61 \times 10^{-6}$ | $4.32 \times 10^{-4} \pm 7.66 \times 10^{-5}$ |

### Purification and quantification of endotoxins

After removal of endotoxins with the Endrotrap HD, the adapted PTMPs were quantified to determine the final concentration of each PTMPs. The CAC_Kpn1_ad PTMP yielded 3.3 x 10^8^ PFU/mL, the CAC_Kpn2_ad PTMP targeting strain B yielded 1.5 x 10^7^ PFU/mL and the CAC_Kpn2_ad PTMPs targeting strain C yielded 1.36 x 10^7^ PFU/mL. The concentration of endotoxin was then quantified using the Pierce Chromatogenic Endotoxin Quant Kit. A calibration curve was first constructed, revealing a linear equation of Y=0.9948X-0.0564 (R2 = 0.9615). Comparison with the calibration curve showed that the CAC_Kpn1_ad, CAC_Kpn2_ad targeting the strain B and CAC_Kpn2_ad targeting the strain C have a value of 0.33 EU/mL, 0.813 EU/mL and 0.813 EU/mL of endotoxin, respectively.

### *G. mellonella* infection model

Determination of the LD_50_ demonstrated that the optimal concentration of *K. pneumoniae* (strains A, B and C) in the inoculum used to infect *G. mellonella* larvae was 10^9^ CFU/mL (data not shown). The larval survival assay showed that the CAC_Kpn1_ad PTMP provided significant protection (p-value < 0.0001) against bacterial infection during 24 h (Figure 5 A). However, for the CAC_Kpn2_ad PTMPs and strains B (Figure 5 B) and C (Figure 5 C), no significant differences were observed between the groups (p-value > 0.05). Both the adapted PTMPs and meropenem alone did not have toxic effects in the larvae (p-value < 0.05).

**Figure 5.**
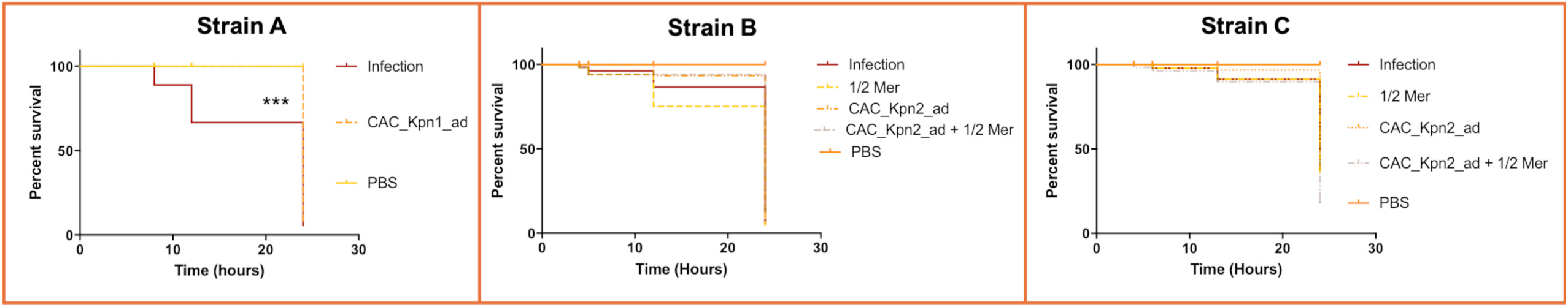
*In vivo* assay with the *G. mellonella* infection model showing the percentage survival at 24 h in the presence of the different adapted PTMPs and the different strains of *K. pneumoniae*: **(A)** A, **(B)** B and **(C)** C.

## DISCUSSION

The increase in emergence of antimicrobial resistance has directed attention towards the use of alternative antimicrobial treatments, such as phages (26). Phage therapy involves the use of phage therapy medical products (PTMPs) to treat infections (26). In this study, two PTMPs, CAC_Kpn1 and CAC_Kpn2, each composed of PTMPs of three seed of phages were designed to target the high-risk *K. pneumoniae* strain ST512. PTMPs are often generated in order to improve the efficacy of phage therapy against bacterial strains and reduce the emergence of phage resistance, an existing problem leading to treatment failure (27–31). In the present study, the PTMPs generated improved the efficacy of the phages against *K. pneumoniae* strains. Notably, no resistance emerged to the CAC_Kpn1 PTMP (composed of the seed of phages vB_KpnM-VAC13, vB_KpnM-VAC36 and vB_KpnM-VAC66) nor CAC_Kpn1_ad using the Appelmans protocol which completely eradicated bacterial growth (8). In the case of the CAC_Kpn2_ad PTMP, (composed by the seed of phages vB_KpnP-VAC25, vB_KpnM-VAC36 and vB_KpnM-VAC66), a great improvement in efficacy was observed relative to the individual phages, but bacterial growth was not completely eradicated. However, there was no great improvement between the adapted and non-adapted PTMPs, and resistance emerged. Thus, beta-lactam meropenem (½ of the MIC), which inhibits wall bacteria synthesis (32), was combined with the CAC_Kpn2_ad PTPM with the aim of reducing the resistance. The combined treatment reduced the emergence of resistance, eradicating the growth of the bacteria up to 20 h in the case of strain B and up to 24 h in the case of strain C. The combination of a sublethal concentration of a ß-lactam antibiotic (½ MIC of meropenem) and PTMPs (CAC_Kpn2_ad and CAC_Kpn2) yielded PAS, considered to have occurred when there is a decrease ≤ 2 -log10 in the cell count of the combination relative to each individual agent (33). According to the CFU/mL counts for strains B and C with the combination of the CAC_Kpn2_ad PTMP together with ½ of the MIC of meropenem, PAS occurred, as there was a reduction of more than 2 -log10 at 12 h and 22 h relative to each antimicrobial agent. This phenomenon has also been observed in other studies. For example, combining meropenem with ciprofloxacin yielded PAS and was effective in controlling *Pseudomonas aeruginosa* and *Candida*. (14) PAS was also observed with a PTMP consisting of phages and antibiotic to eradicate an infection with *Staphyloccocus aureus* in biofilm (12), and PAS led to control of an extensively drug-resistant *Pseudomonas* infection, enabling liver transplantation in an infant (13). In another example, growth of uropathogenic *E. coli* strains was controlled using specifically designed PTMPs, which increased the lytic activity of the phages relative to individual seed of phages, so that 86.7 % of *E. coli* clinical isolates were lysed relative to 50-60 % range of lysis produced by the individual seed of phages. Testing for PAS showed that only seed of phages PS6 and FS17 yielded synergistic effects with the antibiotics amikacin and fosfomycin, and therefore only these seed of phages were included in a PTMP. Combining the PTMP with the antibiotics led to efficient inhibition of bacterial growth in the presence of sublethal doses of antibiotics (34). In another example, combining seed of phages SSE1 and SGF2 with cefotaxime and cefoxinin yielded PAS that halted growth of different strains of *Shigella* (35). More recently, it was demonstrated that a PTMP (HFC-Pae10) composed of three seed of phages specific to *P. aeruginosa* (fNenPA2p2, fNenPA2p4 and fGstaPae02) in combination with meropenem prevented growth of the bacterial more efficiently than phages or antibiotic alone, clearly indicating a synergetic effect (36). Other studies have determined mutant frequencies. Very low mutant frequencies have been observed for strain A in the presence of CAC_Kpn1_ad, with 2.19 x 10^-7^± 9.15 x 10^-8^ mutations per cell. This is consistent with the expected mutant frequency for phage therapy using multiple phage combinations (37, 38), as the combination of multiple phages is believed to limit the emergence of resistance as a wide range of bacterial genotypes are targeted. It has been suggested that resistance to multiple seed of phages may require multiple resistance mutations to arise in the same bacterial cell, with a very low probability of occurrence (39). However, higher mutation frequency rates have been observed with CAC_Kpn2_ad, as indicated by the OD_600nm_ curves and CFU/mL counts. Combining the antibiotic with the CAC_Kpn2_ad PTMP yielded a significantly lower mutation rate, as expected; however, the fact that the mutant frequency in strain C in the presence of the CAC_Kpn2_ad in combination with meropenem was not significantly different from that produced by meropenem alone is surprising. After studying the killing capacity of the PTMPs, the endotoxins were removed from the PTMPs with a view to using the products in animals. The FDA has established a limit of 0.5 EU/mL or 5 EU/mL/kg/ in products designed for therapeutic purposes (40). Therefore, following the elimination of endotoxins, an *in vivo* assay was performed with a *G. mellonella* infection model under the same conditions as the *in vitro* assay. The model was used to observe the effect of the treatment with the CAC_Kpn1_ad and CAC_Kpn2_ad PTMPs on larval survival. The LD_50_ of the inoculum of bacteria was 10^9^ CFUs/mL in all strains, which is 1-Log or 2-Log higher than the inoculum of *K. pneumoniae* in other studies, respectively (41, 42), probably due to the low fitness of these strains. The results obtained in this study indicate an increase in larval survival in the case of the CAC_Kpn1_ad in the strain A during 24 h. In the same line, an increase in larval survival after 4 h of prophylactic treatment and co-injection in three strains was reported (42). However, in case of the post-infection treatment an increase in survival was only observed in one of the strains under study. The same was observed in the present study, where the effect of the CAC_Kpn2_ad PTMP combined with ½ MIC of meropenem was not significantly different from that of the PTMP alone, the ½ MIC of meropenem alone or the adapted PTMPs with ½ MIC of meropenem. In addition, an example of the success of PTMPs *in vivo*, was reported in a study in which the larval survival rate was increased from 55-58 % after 72 h with treatment using individual seed of phages or PTMPs (43). Another study evaluating the application of phages in *G. mellonella* in two different ways, as prophylactic or treatments, against infection by methicillin-resistant *Staphyloccus aureus*. The treatment, consisting of the phage formulation or vancomycin (10 mg/kg), was administered 1 h after the establishment of acute infection. Treatments with different doses of phage or antibiotic administered after inoculation with MRSA improved larval survival. In addition, the PTMP yielded a higher survival rate than the phages alone (44). Finally, a PTMP composed of eight phages was used to treat a 1 h *K. pneumoniae* infection in a *G. mellonella* larvae infection model (41). Following the treatment, the larval survival rates increased from 0 % to 90 % and from 3.3 % to 76.6 % for larvae infected with strains P20 and Kpn2, respectively.

## CONCLUSION

In summary, the preparation of the CAC_Kpn1 and CAC_Kpn2 PTMPs and their subsequent adaptation (CAC_Kpn1_ad and CAC_Kpn2_ad) may be a good approach to solving part of the antimicrobial resistance emergency, as well as the PAS with CAC_Kpn2_ad and ß-lactam antibiotic, allowing the application of phage therapy as a personalized medicine. Moreover, the *in vitro* results were confirmed in the *G. mellonella* models which could be useful in the development of PTMPs against clinical strains of *K. pneumoniae* MDR belonging to clones.

## MATERIAL AND METHODS

### In vitro assay

The workflow of the *in vitro* assay is summarized in Figure 6.

**Figure 6.**
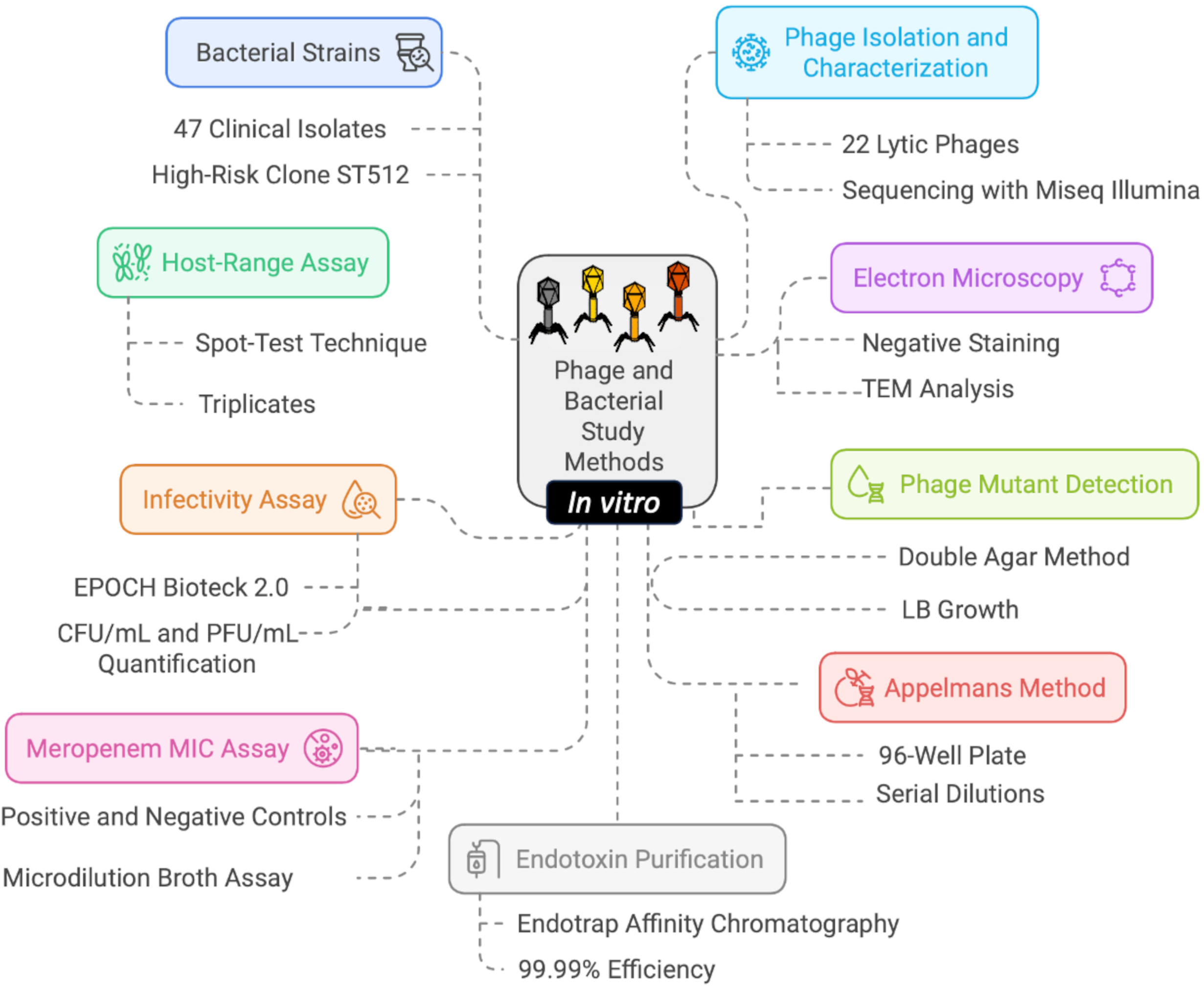
*In vitro* assay workflow. The workflow was performed with the Napkin software.

### Bacterial strains

A collection of 47 previously identified clinical isolates of *K. pneumoniae* obtained from the Virgen Macarena University Hospital (Seville, Spain) and the National Center of Microbiology (Carlos III Health Institute, Spain) (45) was used in the present study. In addition, three clinical strains of the high-risk ST512 clone producing KPC-3 carbapenemase were used: A, B and C. These strains come from different Spanish geographical regions and had genetic distances between them greater than 20 alleles (21–34), using a core genome MLST (cgMLST) scheme consisting of 2,538 *K. pneumoniae* targets provided by SeqSphere version 3.5.0 (Ridom, Münsten, Germany).

### Isolation, purification, propagation and sequencing of seed of phages

Twenty-two lytic phages from MePRAM collection were used to produce PTMPs. The process of isolating, purifying, propagating, characterizing and sequencing the seed of phages with the MiSeq System (Illumina) has already been described (45). All genomes are available in the GenBank Umbrella Bioproject PRJNA1101025 and the Bioproject PRNJA739095.

### Host-range assay

The phage host-range was determined by the spot-test technique (46), in the clinical strains of *K. pneumoniae* of the high-risk ST512 clone isolated in the previously mentioned study (45). Briefly, a negative control consisting of SM buffer (0.1 NaCl, 10 mM MgSO_4_, 20 mM Tris-HCL; pH 7.5) was included in each plate. Phage infectivity was established by the presence of clear spots (infection), the presence of turbid spots (low infection or resistance) and the lack of spots (no infection). All determinations were carried out in triplicate. Once the seed of phages were selected for generating the PTMPs, comparative genomic analysis was conducted using the VipTree server (https://www.genome.jp/viptree/).

### Electron microscopic analysis of seed of phages

The four seed of lytic phages selected for generating the PTMPs were negatively stained with 1 % aqueous uranyl acetate before being analyzed by (TEM) electron microscopy (JEOL JEM-1011).

### Infectivity assay in liquid media

The phage infectivity assay was performed in liquid media by measuring the absorbance as the optical density at a wavelength of 600 nm (OD_600nm_) every 15 min for 22 h with shaking at 540 rpm in a microplate reader (EPOCH Bioteck 2.0, Agilent). The experiment was conducted with the seed of phages selected in the spot-test assay (vB_KpnM-VAC13, vB_KpnP-VAC25, vB_KpnM-VAC36, vB_KpnM-VAC66) and with the not-adapted (CAC_Kpn1) and adapted (CAC_Kpn1_ad) PTMPs (vB_KpnM-VAC13, vB_KpnM-VAC36 and vB_KpnM-VAC66) and with the not adapted (CAC_Kpn2) and adapted (CAC_Kpn2_ad) PTMPs (vB_KpnP-VAC25, vB_KpnM-VAC36 and vB_KpnM-VAC66). In the case of the CAC_Kpn2_ad PTMP, the infectivity assay was also performed with half the minimum inhibitory concentration (MIC) of meropenem. Finally, the number of colony forming units (CFUs)/mL and phage forming units (PFUs)/mL were quantified at 12 h and 22 h. All experiments were performed in triplicate.

### The Appelmans protocol

The PTMPs produced (CAC_Kpn1 and CAC_Kpn2) were adapted using the Appelmans protocol (6, 10). Briefly, 180 µL of Luria-Bertani (LB) broth was added to the wells of microtitre plates (96-wells) along with 20 µL of PTMP. Serial dilutions of the PTMP were then performed. Finally, 20 µL of bacteria (1 x 10^6^ CFU/mL) was added to each well and the plates were incubated at 37°C for 24 h with shaking at 180 rpm. The contents of the wells in which lysate was observed were then collected. If no lysis was observed, the contents of the first well were collected, i.e. the well with the highest concentration of seed of phages. Chloroform (1 %) was added to the samples collected and incubated for 15 to 20 min in cold incubation. The samples were then centrifuged at 14,000 x g for 15 min and the supernatant was used as new stock for preparing a new plate. The adaptation process was continued over a maximum of 5 days.

### Meropenem minimal inhibitory concentration assay

The meropenem MIC for *K. pneumoniae* clinical strains B and C was established by microdilution broth assay (15). Briefly, eleven serial double dilutions of meropenem were prepared in Muller-Hinton broth (MHB) in 96-well microtiter plates. Each well was then inoculated with the corresponding *K. pneumoniae* strain to a final concentration of 5 x 10^5^ CFU/mL, diluted from an overnight culture. A row of MHB inoculated with *K. pneumoniae* was included as a positive control and a row with only MHB was included as a negative control. The plate was incubated for 24 h at 37 °C, and the MIC was subsequently determined as the concentration of meropenem in the first well where no bacterial growth was observed (15). All experiments were performed in triplicate.

### Determination of the rate of emergence of phage-resistant bacterial mutants

The frequency of occurrence of phage-resistant mutant bacterial cells was detected by the method described in Merabishvili et *al.* (47). Briefly, five isolated colonies were selected and inoculated into five tubes with 4 mL of LB, which were incubated at 37°C overnight at 180 rpm. Aliquots 100 µL of 10^0^ to 10^-4^ dilutions of bacterial culture at 10^8^ CFU/mL were brought into contact with 100 µL aliquots of the 10^9^ PFU/mL phage broth, and the total volume of 200 µL was inoculated onto TA plates by the double agar method and incubated at 37 °C for several days. Simultaneously, 100 µL aliquots of bacterial culture diluted by 10^-5^ to 10^-6^ were plated on LB medium. The frequency of mutant cells was calculated by dividing the number of resistant bacteria by the total number of sensitive bacteria. The frequency of occurrence of antibiotic-phage mutant bacteria in response to the meropenem and PTMP combination was then determined, as previously described but with the addition of ½ MIC of the antibiotic to the medium.

### Purification of endotoxins

Endotoxins were purified on an affinity chromatography column (EndoTrap®HD, Hyglos, Bernried am Stranberger Seen, Germany), in which a bacteriophage-derived protein in the column matrix binds endotoxins with high affinity and specificity. According to the manufacturer, EndoTrap®HD kit can remove endotoxins from proteins, peptides, antibodies, RNA/DNA, antigens and plant extracts, with a removal efficiency of 99.99%. The adapted CAC_Kpn1 and CAC_Kpn2 PTPMs were separated from endotoxins using the commercial EndoTrap®HD kit, according to the manufacturer’s instructions. Briefly, the PTPMs were diluted 1/10 in phosphate buffer saline (PBS) supplemented with 1 mM CaCl_2_ to an estimated final concentration of 10^8^ PFU/mL.

### Quantification of endotoxins

The amount of endotoxin in the adapted CAC_Kpn1_ad and CAC_Kpn2_ad PTMPs was quantified using the Pierce Chromatogenic Endotoxin Quant Kit (Thermo Fisher). The kit is based on the amoebocyte lysate assay, which has become the standard for endotoxin detection. The kit detects endotoxins, measured as endotoxin units (EU) per millilitre, at levels as low as 0.01 EU/mL. The PTMPs were quantified using the double agar method, to enable calculation of the actual number of PFUs/mL.

### In vivo assay

The workflow of the *in vivo* assay with the *Galleria mellonella* model is outlined in Figure 7.

**Figure 7.**
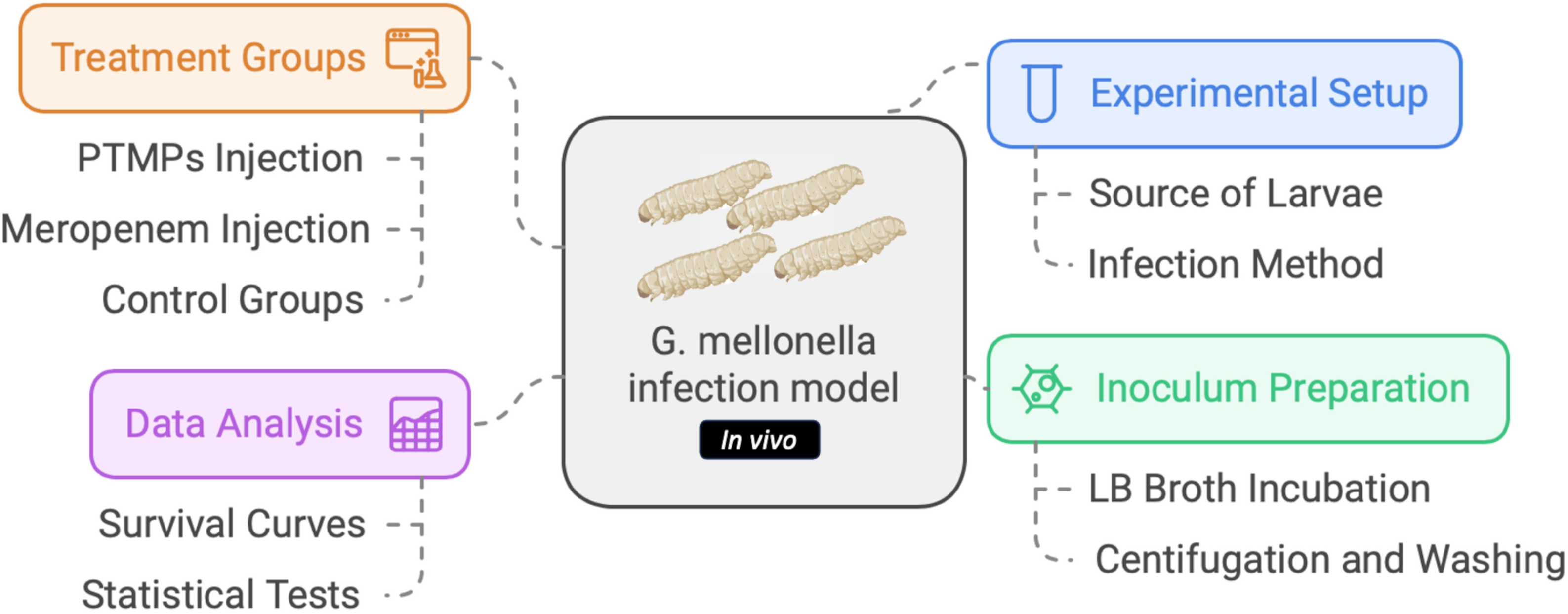
*In vivo* assay workflow. The workflow was performed with the Napkin software.

### *G. mellonella* infection model

The *G. mellonella* model was adapted from the version developed by Peleg et *al.* (48) and Blasco and *al*. (49). Briefly, fifteen *G. mellonella* larvae, obtained from a local pet shop (Harkito, Madrid), were infected by injection of 10^9^ CFU/mL of adapted PTMPs (diluted in 10 µL of PBS) in the last left proleg, with a Hamilton syringe (Hamilton, Shanghai, China). The concentration of inoculum was selected after determination of the 50 % lethal dose (LD_50_), i.e. the amount of inoculum that causes death of 50 % of the larvae 24 h post-infection (49). For preparation of the inoculum, 5 mL of LB broth was incubated with the bacteria overnight at 37°C with shaking at 180 rpm. The culture was then centrifuged and separated by centrifugation (4,000 x g for 15 min) and washed twice with PBS. One hour after being injected with the bacteria, the larvae were injected with 10 µL of one of the following treatments: PTMP (CAC_Kpn1_ad or CAC_Kpn2_ad) at a concentration of 10^9^ PFU/mL (previously purified by EndoTrap®HD kit and concentrated by Amikon); meropenem (½ MIC); CAC_Kpn2_ad plus meropenem (½ MIC); or adapted PTMPs and meropenem (as control). All treatments were administered at the same concentrations used in the *in vitro* assay. A control group of infected larvae was injected with PBS alone (no treatment). The treated larvae were placed in Petri dishes and incubated in darkness at 37°C. The number of dead larvae was recorded after 24 h. The larvae were considered dead when they showed no movement in response to touch. The survival curves were plotted using GraphPad Prism v.6, and data were analysed using the Gehan-Breslow-Wilcoxon test.

